# PathCORE-T: identifying and visualizing globally co-occurring pathways in large transcriptomic compendia

**DOI:** 10.1101/147645

**Authors:** Kathleen M. Chen, Jie Tan, Gregory P. Way, Georgia Doing, Deborah A. Hogan, Casey S. Greene

## Abstract

**Background:** Investigators often interpret genome-wide data by analyzing the expression levels of genes within pathways. While this within-pathway analysis is routine, the products of any one pathway can affect the activity of other pathways. Past efforts to identify relationships between biological processes have evaluated overlap in knowledge bases or evaluated changes that occur after specific treatments. Individual experiments can highlight condition-specific pathway-pathway relationships; however, constructing a complete network of such relationships across many conditions requires analyzing results from many studies.

**Results:** We developed PathCORE-T framework by implementing existing methods to identify pathway-pathway transcriptional relationships evident across a broad data compendium. PathCORE-T is applied to the output of feature construction algorithms; it identifies pairs of pathways observed in features more than expected by chance as *functionally co-occurring*. We demonstrate PathCORE-T by analyzing an existing eADAGE model of a microbial compendium and building and analyzing NMF features from the TCGA dataset of 33 cancer types. The PathCORE-T framework includes a demonstration web interface, with source code, that users can launch to (1) visualize the network and (2) review the expression levels of associated genes in the original data. PathCORE-T creates and displays the network of globally co-occurring pathways based on features observed in a machine learning analysis of gene expression data.

**Conclusions:** The PathCORE-T framework identifies transcriptionally co-occurring pathways from the results of unsupervised analysis of gene expression data and visualizes the relationships between pathways as a network. PathCORE-T recapitulated previously described pathway-pathway relationships and suggested experimentally testable additional hypotheses that remain to be explored.

## Background

The number of publicly available genome-wide datasets is growing rapidly [1]. High-throughput sequencing technologies that measure gene expression quickly with high accuracy and low cost continue to enable this growth [2]. Expanding public data repositories [3, 4] have laid the foundation for computational methods that consider entire compendia of gene expression data to extract biological patterns [5]. These patterns may be difficult to detect in measurements from a single experiment. Unsupervised approaches, which identify important signals in the data without being constrained to previously-described patterns, may discover new expression modules and thus will complement supervised methods, particularly for exploratory analyses [6,7].

Feature extraction methods are a class of unsupervised algorithms that can reveal unannotated biological processes from genomic data [7]. Each feature can be defined by a subset of influential genes, and these genes suggest the biological or technical pattern captured by the feature. These features, like pathways, are often considered individually [7,8]. When examined in the context of knowledgebases such as the Kyoto Encyclopedia of Genes and Genomes (KEGG) [9], most features are significantly enriched for more than one biological gene set [7]. In this work, we refer to such a gene set by the colloquial term, pathway. It follows then that such features can be described by sets of functionally related pathways. We introduce the PathCORE-T (identifying ***path***way ***co***-occurrence ***re***lationships in ***t***ranscriptomic data) software, which implements existing methods that jointly consider features and gene sets to map pathways with shared transcriptional responses.

PathCORE-T offers a data-driven approach for identifying and visualizing transcriptional pathway-pathway relationships. In this case, relationships are drawn based on the sets of pathways, annotated in a resource of gene sets, occurring within constructed features. Because PathCORE-T starts from a feature extraction model, the number of samples in the compendium used for model generation and the fraction of samples needed to observe a specific biological or technical pattern is expected to vary by feature extraction method. Pathways must be perturbed in a sufficient fraction of experiments in the data compendium to be captured by any such method. To avoid discovering relationships between pathways that share many genes—which could more easily be discovered by directly comparing pathway membership—we implement an optional pre-processing step that corrects for genes shared between gene sets, which Donato et al. refer to as pathway crosstalk [10]. Donato et al.’s correction method, maximum impact estimation, has not previously been implemented in open source software. We have released our implementation of maximum impact estimation as its own Python package (PyPI package name: crosstalk-correction) so that it can be used independently of PathCORE-T. Applying this correction in PathCORE-T software allows a user to examine relationships between gene sets based on how genes are expressed as opposed to which genes are shared.

We apply PathCORE-T to a microbial and a cancer expression dataset, each analyzed using different feature extraction methods, to demonstrate its broad applicability. For the microbial analysis, we created a network of KEGG pathways from recently described ensemble Analysis using Denoising Autoencoders for Gene Expression (eADAGE) models trained on a compendium of Pseudomonas aeruginosa (*P. aeruginosa*) gene expression data (doi:10.5281/zenodo.583694) [7]. We provide a live demo of the PathCORE-T web application for this network: users can click on edges in the network to review the expression levels of associated genes in the original compendium (https://pathcore-demo.herokuapp.com/PAO1). To show its use outside of the microbial space, we also demonstrate PathCORE-T analysis of Pathway Interaction Database (PID)-annotated [11] non-negative matrix factorization (NMF) features [12, 13] extracted from The Cancer Genome Atlas’s (TCGA) pan-cancer dataset of 33 different tumor types (doi:10.5281/zenodo.56735) [14].

In addition to visualizing the results of these two applications, the PathCORE-T web interface (https://pathcore-demo.herokuapp.com/) links to the documentation and source code for our implementation and example usage of PathCORE-T. Methods implemented in PathCORE-T are written in Python and pip-installable (PyPI package name: PathCORE-T). Examples of how to use these methods are provided in the PathCORE-T analysis repository (https://github.com/greenelab/PathCORE-T-analysis). In addition to scripts that reproduce the eADAGE and NMF analyses described in this paper, the PathCORE-T-analysis repository includes a Jupyter notebook (https://goo.gl/VuzN12) with step-by-step descriptions for the complete PathCORE-T framework.

### Related work

Our approach diverges from other algorithms that we identified in the literature in its intent: PathCORE-T finds pathway pairs within a biological system that are overrepresented in features constructed from diverse transcriptomic data. This complements other work that developed models specific to a single condition or disease. Approaches designed to capture pathway-pathway interactions from gene expression experiments for disease-specific, case-control studies have been published [15,16]. For example, Pham et al. developed Latent Pathway Identification Analysis to find pathways that exert latent influences on transcriptionally altered genes [17]. Under this approach, the transcriptional response profiles for a binary condition (disease/normal), in conjunction with pathways specified in the KEGG and functions in Gene Ontology (GO) [18], are used to construct a pathway-pathway network where key pathways are identified by their network centrality scores [17]. Similarly, Pan et al. measured the betweenness centrality of pathways in disease-specific genetic interaction and coexpression networks to identify those most likely to be associated with bladder cancer risk [19]. These methods captured pathway relationships associated with a particular disease state.

Global networks identify relationships between pathways that are not disease- or condition-specific. One such network, detailed by Li et al., relied on publicly available protein interaction data to determine pathway-pathway interactions [20]. Two pathways were connected in the network if the number of protein interactions between the pair was significant with respect to the computed background distribution. Such approaches rely on databases of interactions, though the interactions identified can be subsequently used for pathway-centric analyses of transcriptomic data [20, 21]. Pita-Juárez et al. created the Pathway Coexpression Network (PCxN) as a tool to discover pathways correlated with a pathway of interest [22]. They estimated correlations between pathways based on the expression of their underlying genes (as annotated in MSigDB) across a curated compendium of microarray data [22]. Software like PathCORE-T that generates global networks of pathway relationships from unsupervised feature analysis models built using transcriptomics data has not yet been published.

The intention of PathCORE-T is to work from transcriptomic data in ways that do not give undue preference to combinations of pathways that share genes. Other methods have sought to consider shared genes between gene sets, protein-protein interactions, or other curated knowledgebases to define pathway-pathway interactions [20–21, 23–25]. For example, Glass and Girvan described another network structure that relates functional terms in GO based on shared gene annotations [26]. In contrast with this approach, PathCORE-T specifically removes gene overlap in pathway definitions before they are used to build a network. Our software reports pathway-pathway connections overrepresented in gene expression patterns extracted from a large transcriptomic compendium while controlling for the fact that some pathways share genes.

## Implementation

PathCORE-T identifies functional links between known pathways from the output of feature construction methods applied to gene expression data (Fig. 1a, b). The result is a network of pathway co-occurrence relationships that represents the grouping of biological gene sets within those features. We correct for gene overlap in the pathway annotations to avoid identifying co-occurrence relationships driven by shared genes. Additionally, PathCORE-T implements a permutation test for evaluating and removing edges—pathway-pathway relationships—in the resulting network that cannot be distinguished from a null model of random associations. Though we refer to the relationships in the network as co-occurrences, it is important to note that the final network displays co-occurrences that have been filtered based on this permutation test (Fig. 1c).

**Figure 1.**
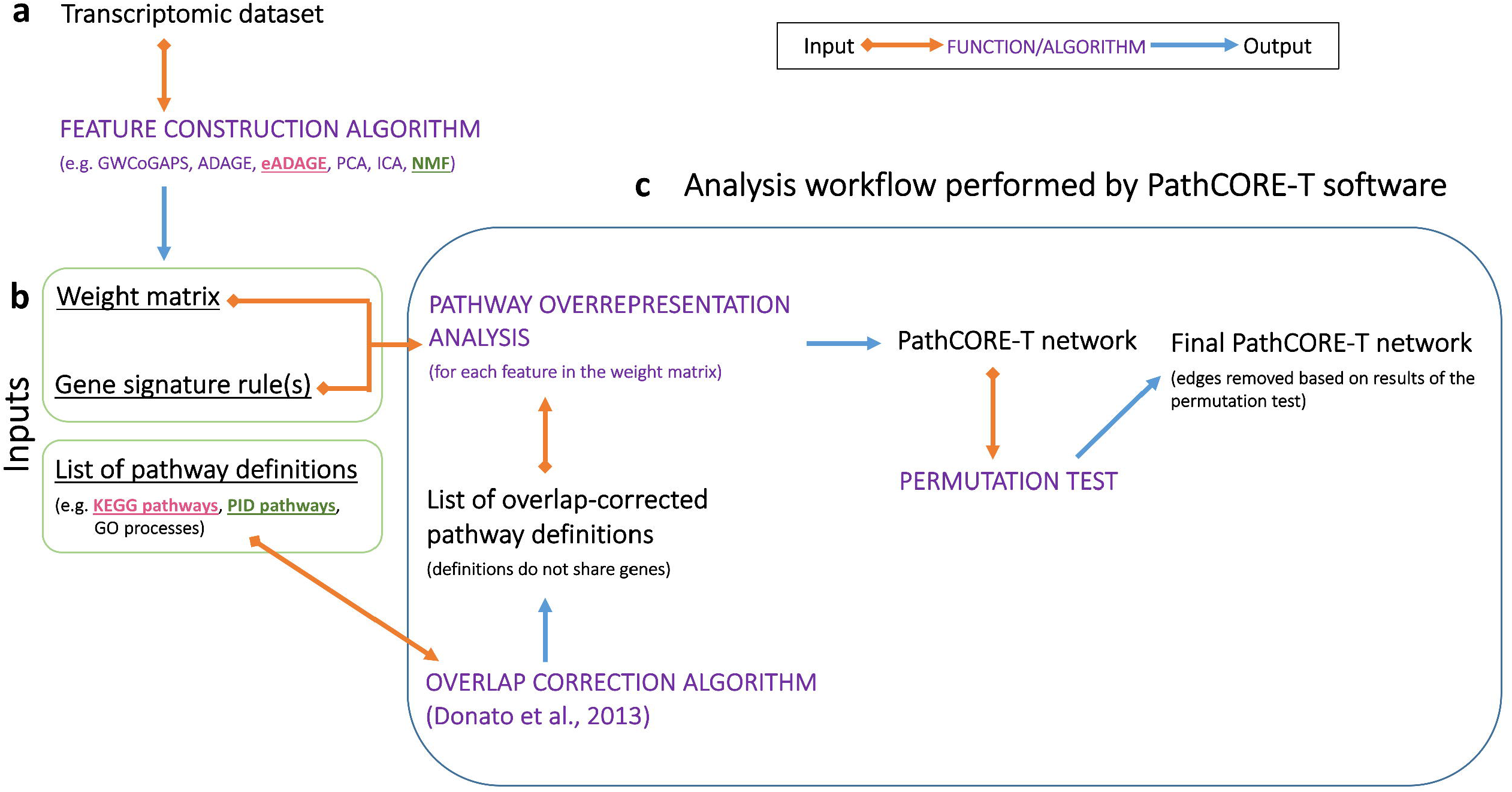
The PathCORE-T software analysis workflow. **(a)** A user applies a feature construction algorithm to a transcriptomic dataset of genes-by-samples. The features constructed must preserve the genes in the dataset and assign weights to each of these genes according to some distribution. **(b)** Inputs required to run the complete PathCORE-T analysis workflow. The constructed features are stored in a weight matrix. Based on how gene weights are distributed in the constructed features, a user defines thresholds to select the set of genes most indicative of each feature’s function—we refer to these user-defined thresholds as gene signature rules. Finally, a list of pathway definitions will be used to interpret the features and build a pathway co-occurrence network. **(c)** Methods in the PathCORE-T analysis workflow (indicated using purple font) can be employed independently of each other so long as the necessary input(s) are provided. The 2 examples we describe to demonstrate PathCORE-T software use the following inputs: (1) the weight matrix and gene signature rules for eADAGE (applied to the *P. aeruginosa* gene compendium) and KEGG pathways and (2) the weight matrix and gene signature rules for NMF (applied to the TCGA pan-cancer dataset) and PID pathways.

Our software is written in Python and pip-installable (PyPI package name: PathCORE-T), and examples of how to use the methods in PathCORE-T are provided in the PathCORE-T-analysis repository (https://github.com/greenelab/PathCORE-T-analysis). We recommend that those interested in using the PathCORE-T software consult the documentation and scripts in PathCORE-T-analysis. Each of the functions in PathCORE-T that we describe here can be used independently; however, we expect most users to employ the complete approach for interpreting pathways shared in extracted features (Fig. 1).

### Data organization

PathCORE-T requires the following inputs:

1. **A weight matrix** that connects each gene to each feature. We expect that this results from the application of a feature construction algorithm to a compendium of gene expression data. The primary requirements are that features must contain the full set of genes in the compendium and genes must have been assigned weights that quantify their contribution to a given feature. Accordingly, a weight matrix will have the dimensions *n* × *k*, where *n* is the number of genes in the compendium and *k* is the number of features constructed. In principal component analysis (PCA), this is the loadings matrix [27]; in independent component analysis (ICA), it is the unmixing matrix [28]; in ADAGE or eADAGE it is termed the weight matrix [5,7]; in NMF it is the matrix *W*, where the NMF approximation of the input dataset *A* is *A* ~ *WH* [12]. In addition to the scripts we provide for the eADAGE and NMF examples in the PathCORE-T analysis repository, we include a Jupyter notebook (https://goo.gl/VuzN12) that demonstrates how a weight matrix can be constructed by applying ICA to the *P. aeruginosa* gene compendium.
2. **Gene signature rule(s).** To construct a pathway co-occurrence network, the weight matrix must be processed into gene signatures by applying threshold(s) to the gene weights in each feature—we refer to these as gene signature rules. Subsequent pathway overrepresentation will be determined by the set of genes that makes up a feature’s gene signature. These are often the weights at the extremes of the distribution. How gene weights are distributed will depend on the user’s selected feature construction algorithm; because of this, a user must specify criterion for including a gene in a gene signature. PathCORE-T permits rules for a single gene signature or both a positive and a negative gene signature. The use of 2 signatures may be appropriate when the feature construction algorithm produces positive and negative weights, the extremes of which both characterize a feature (e.g. PCA, ICA, ADAGE or eADAGE). Because a feature can have more than one gene signature, we maintain a distinction between a feature and a feature’s gene signature(s).
3. **A list of pathway definitions**, where each pathway is defined by a set of genes (e.g. KEGG pathways, PID pathways, GO biological processes). We provide the files for the *P. aeruginosa* KEGG pathway definitions and the Nature-NCI PID pathway definitions in the PathCORE-T analysis repository (https://github.com/greenelab/PathCORE-T-analysis/tree/master/data).

### Weight matrix construction and signature definition

In practice, users can obtain a weight matrix from many different methods. For the purposes of this paper, we demonstrate generality by constructing weight matrices via eADAGE and NMF.

#### eADAGE

eADAGE is an unsupervised feature construction algorithm developed by Tan et al. [7] that uses an ensemble of neural networks (an ensemble of ADAGE models) to capture biological patterns embedded in the expression compendium. We use models from Tan et al. [7]. In that work, Tan et al. evaluated multiple eADAGE model sizes to identify that *k*=300 features was an appropriate size for the current *P. aeruginosa* compendium. The authors also compared eADAGE to two other commonly used feature construction approaches, PCA and ICA [7]. Tan et al. produced 10 eADAGE models that each extracted *k*=300 features from the compendium of genome-scale *P. aeruginosa* data. Because PathCORE-T supports the aggregation of co-occurrence networks created from different models on the same input data, we use all 10 of these models in the PathCORE-T analysis of eADAGE models (doi:10.5281/zenodo.583172).

Tan et al. refers to the features constructed by eADAGE as nodes. They are represented as a weight matrix of size *n* × *k*, where *n* genes in the compendium are assigned positive or negative gene weights, according to a standard normal distribution, for each of the *k* features. Tan et al. determined that the gene sets contributing the highest positive or highest negative weights (+/− 2.5 standard deviations) to a feature described gene expression patterns across the compendium, and thus referred to the gene sets as signatures. Because a feature’s positive and negative gene signatures did not necessarily correspond to the same biological processes or functions, Tan et al. analyzed each of these sets separately [7]. Tan et al.’s gene signature rules are specified as an input to the PathCORE-T analysis as well.

#### NMF

We also constructed an NMF model for the TCGA pan-cancer dataset. Given an NMF approximation of *A* ~ *WH* [12], where *A* is the input expression dataset of size *n* × *s* (*n* genes by *s* samples), NMF aims to find the optimal reconstruction of *A* by *WH* such that *W* clusters on samples (size *n* × *k*) and *H* clusters on genes (size *k* × *s*). In order to match the number of features constructed in each eADAGE model by Tan et al., we set *k*, the desired number of features, to be 300 and used *W* as the input weight matrix for the PathCORE-T software. We found that the gene weight distribution of an NMF feature is right-skewed and (as the name suggests) non-negative (Fig. S1). In this case, we defined a single gene signature rule: an NMF feature’s gene signature is the set of genes with weights 2.0 standard deviations above the mean weight of the feature.

The selection of *k*=300 for the NMF model allowed us to make the eADAGE-based and NMF-based case studies roughly parallel. We verified that 300 components was appropriate by evaluating the percentage of variance explained by PCA applied to the TCGA dataset. In general, the principal components explained very little variance—the first principal component only explained 11% of the variance. At 300 components, the proportion of variance explained was 81%.

As an additional analysis, we determined the number of components (*k*=24) where each additional component explained less than 0.5% of the variance. We found that using a very small number of constructed features resulted in a substantial loss of power: PathCORE-T analysis with a single *k*=24 model yielded no significant edges after permutation test. However, PathCORE-T can be applied over multiple models as long as the feature construction method produces different solutions depending on random seed initialization. We performed 10 factorizations to generate an aggregate of 10 *k*=24-feature NMF models and found that the resulting co-occurrence network was denser (364 edges) than our *k*=300 factor network (119 edges). 65 edges were found in both networks. These shared edges had higher weights, on average, in both networks compared to edges unique to each network (https://goo.gl/vnDVNA).

### Construction of a pathway co-occurrence network

We employ a Fisher’s exact test [29] to determine the pathways significantly associated with each gene signature. When considering significance of a particular pathway, the two categories of gene classification are as follows: (1) presence or absence of the gene in the gene signature and (2) presence or absence of the gene in the pathway definition. For each pathway in the input list of pathway definitions, we specify a contingency table and calculate its p-value, which is corrected using the Benjamini—Hochberg [30] procedure to produce a feature-wise false discovery rate (FDR). A pathway with an adjusted p-value that is less than the user-settable FDR significance cutoff, alpha (default: 0.05), is considered significantly enriched in a given gene signature. This cutoff value should be selected to most aid user interpretation of the model. The next step of PathCORE-T is to convert pathway-node relationships into pathway-pathway relationships. For this, we apply a subsequent permutation test over pathway-pathway edge weights that accounts for the frequency at which pathways are observed as associated with features. This permutation produces a p-value for each edge. Two pathways co-occur, or share an edge in the pathway co-occurrence network, if they are both overrepresented in a gene signature. The weight of each edge in the pathway-pathway graph corresponds to number of times such a pathway pair is present over all gene signatures in a model (Fig. 2a).

**Figure 2.**
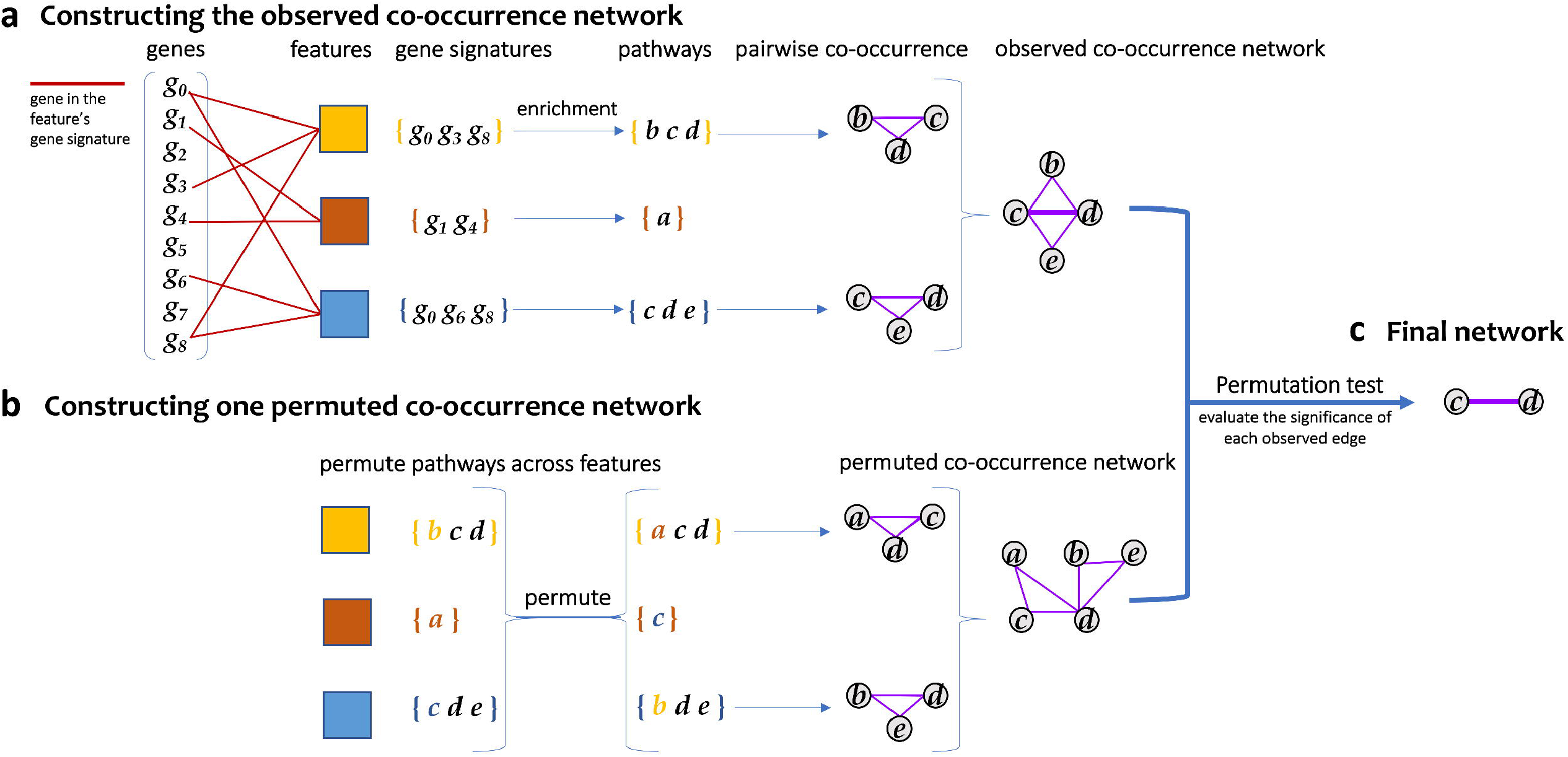
The approach implemented in PathCORE-T to construct a pathway co-occurrence network from an expression compendium. **(a)** A user-selected feature extraction method is applied to expression data. Such methods assign each gene a weight, according to some distribution, that represents the gene’s contribution to the feature. The set of genes that are considered highly representative of a feature’s function is referred to as a feature’s gene signature. The gene signature is user-defined and should be based on the weight distribution produced by the unsupervised method of choice. In the event that the weight distribution contains both positive and negative values, a user can specify criteria for both a positive and negative gene signature. A test of pathway enrichment is applied to identify corresponding sets of pathways from the gene signature(s) in a feature. We consider pathways significantly overrepresented in the same feature to co-occur. Pairwise co-occurrence relationships are used to build a network, where each edge is weighted by the number of features containing both pathways. **(b)** *N* permuted networks are generated to assess the statistical significance of a co-occurrence relation in the graph. Here, we show the construction of one such permuted network. Two invariants are maintained during a permutation: (1) pathway side-specificity (if applicable, e.g. positive and negative gene signatures) and (2) the number of distinct pathways in a feature’s gene signature. **(c)** For each edge observed in the co-occurrence network, we compare its weight against the weight distribution generated from *N* (default: 10,000) permutations of the network to determine each edge’s p-value. After correcting the p-value by the number of edges observed in the graph using the Benjamini—Hochberg procedure, only an edge with an adjusted p-value below alpha (default: 0.05) is kept in the final co-occurrence network.

### Permutation test

The network that results from the preceding method is densely connected, and many edges may be spurious. To remove correlations that cannot be distinguished from random pathway associations, we define a statistical test that determines whether a pathway-pathway relationship appearing *x* times in a *k*-feature model is unexpected under the null hypothesis—the null hypothesis being that the relationship does not appear more often than it would in a random network. We create *N* weighted null networks, where each null network is constructed by permuting overrepresented pathways across the model’s gene signatures while preserving the number of pathways for which each gene signature is enriched (Fig. 2b). *N* is a user-settable parameter: the example PathCORE-T analyses we provide specify an *N* of 10,000. Increasing the value of *N* leads to more precise p-values, particularly for low p-values, but comes at the expense of additional computation time.

In the case where we have positive and negative gene signatures, overrepresentation can be positive or negative. Because certain pathways may display bias toward one side—for example, a pathway may be overrepresented more often in features’ positive gene signatures—we perform the permutation separately for each side. The *N* random networks produce the background weight distribution for every observed edge; significance can then be assessed by comparing the true (observed) edge weight against the null. The p-value for each edge *e* is calculated by summing the number of times a random network contained *e* at a weight greater than or equal to its observed edge weight and dividing this value by *N*. Following Benjamini—Hochberg FDR correction by the number of edges in the observed network, pathway-pathway relationships with adjusted p-values above alpha (user-settable default: 0.05) are removed from the network of co-occurring pathways (Fig. 2c). The threshold alpha value is a configurable parameter, and the user should select an FDR that best balances the costs and consequences of false positives. For highly exploratory analyses in which it may be helpful to have more speculative edges, this value can be raised. For analyses that require particularly stringent control, it can be lowered.

Because the expected weight of every edge can be determined from the *N* random networks (by taking the sum of the background weight distribution for an edge and dividing it by *N*), we can divide each observed edge weight by its expected weight (dividing by 1 if the expected edge weight is 0 based on the *N* permutations) to get the edge’s odds ratio. Edges in the final network are weighted by their odds ratios.

### Gene overlap correction

Pathways can co-occur because of shared genes (Fig. 3a, b, d). Though some approaches use the overlap of genes to identify connected pathways, we sought to capture pairs of pathways that persisted even when this overlap was removed. The phenomenon of observing enrichment of multiple pathways due to gene overlap has been previously termed as “crosstalk,” and Donato et al. have developed a method to correct for it [10]. Due to confusion around the term, we refer to this as overlapping genes in this work, except where specifically referencing Donato et al. Their approach, called maximum impact estimation, begins with a membership matrix indicating the original assignment of multiple genes to multiple pathways. It uses expectation maximization to estimate the pathway in which a gene contributes its greatest predicted impact (its maximum impact) and assigns the gene only to this pathway [10]. This provides a set of new pathway definitions that no longer share genes (Fig. 3c, e).

**Figure 3.**
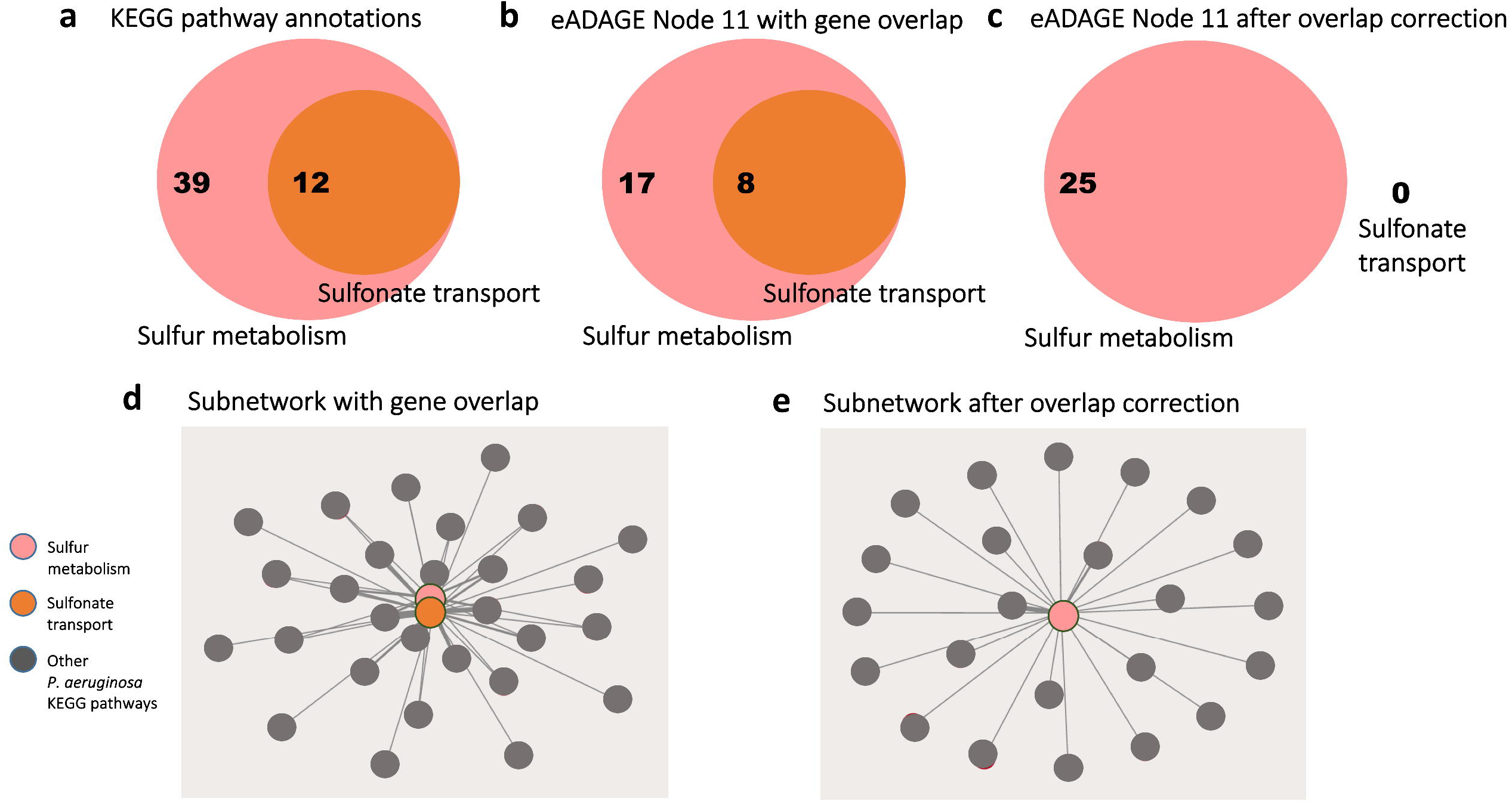
Correcting for gene overlap results in a sparser pathway co-occurrence network. **(a)** The KEGG pathway annotations for the sulfonate transport system are a subset of those for sulfur metabolism. 12 genes annotated to the sulfonate transport system are also annotated to sulfur metabolism. **(b)** Without applying the overlap correction procedure, 25 of the genes in the positive and negative gene signatures of the eADAGE feature “Node 11” are annotated to sulfur metabolism--of those, 8 genes are annotated to the sulfonate transport system as well. **(c)** All 8 of the overlapping genes are mapped to the sulfur metabolism pathway after overlap correction. **(d)** A co-occurrence network built without applying the overlap correction procedure will report co-occurrence between the sulfonate transport system and sulfur metabolism, whereas **(e)** no such relation is identified after overlap correction.

There was no existing open source implementation of this algorithm, so we implemented Donato et al.’s maximum impact estimation as a Python package (PyPI package name: crosstalk-correction). This software is separate from PathCORE-T because we expect that it may be useful in its own right for other analytical workflows, such as differential expression analysis. The procedure is written using NumPy functions and data structures, which allows for efficient implementation of array and matrix operations in Python [31].

In PathCORE-T, we used this software to resolve overlapping genes before pathway overrepresentation analysis. Overlap correction is applied to each feature of the model independently. This most closely matches the setting evaluated by the original authors of the method. Work on methods that resolves overlap by using information shared across features may provide opportunities for future enhancements but was deemed to be out of the scope of a software contribution.

With this step, the pathway co-occurrence network identifies relationships that are not driven simply by the same genes being annotated to multiple pathways. Without this correction step, it is difficult to determine whether a co-occurrence relationship can be attributed to the features extracted from expression data or gene overlap in the two pathway annotations. We incorporate this correction into the PathCORE-T workflow by default; however, users interested in using PathCORE-T to find connections between overlapping gene sets can choose to disable the correction step.

### PathCORE-T network visualization and support for experimental follow-up

The PathCORE-T analysis workflow outputs a list of pathway-pathway relationships, or edges in a network visualization, as a text file. An example of the KEGG *P. aeruginosa* edges file is available for download on the demo application: http://pathcore-demo.herokuapp.com/quickview. While we chose to represent pathway-pathway relationships as a network, users can use this file output to visualize the identified relationships as an adjacency matrix or in any other format they choose.

As an optional step, users can set up a Flask application for each PathCORE-T network. Metadata gathered from the analysis are saved to TSV files, and we use a script to populate collections in a MongoDB database with this information. The co-occurrence network is rendered using the D3.js force-directed graph layout [32]. Users can select a pathway-pathway relationship in the network to view a new page containing details about the genes annotated to one or both pathways (Fig. 4a).

**Figure 4.**
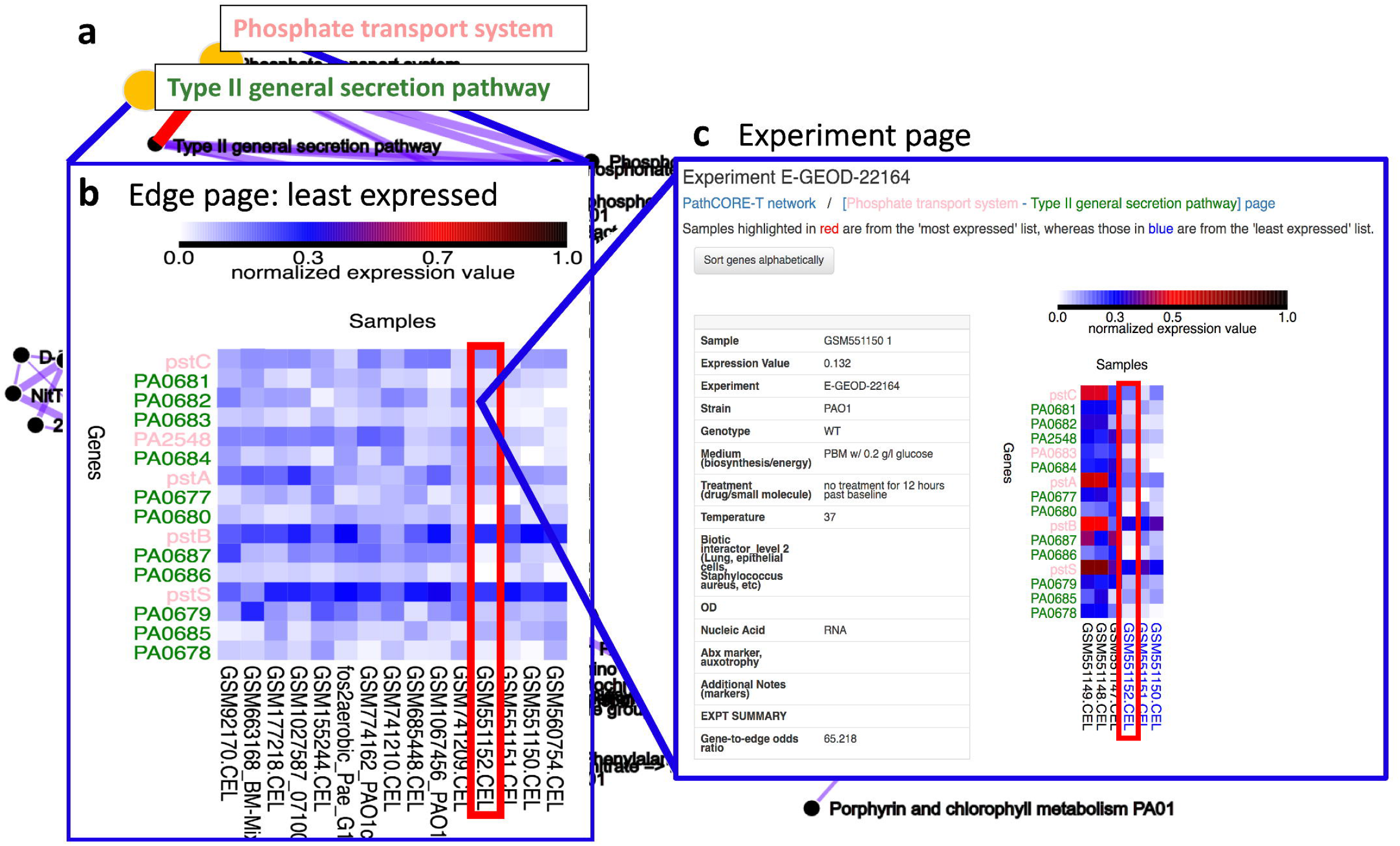
A web application used to analyze pathway-pathway relationships in the eADAGE-based, *P. aeruginosa* KEGG network. **(a)** A user clicks on an edge (a pathway-pathway relationship) in the network visualization. (b) The user is directed to a page that displays expression data from the original transcriptomic dataset specific to the selected edge (https://goo.gl/Hs5A3e). The expression data is visualized as two heatmaps that indicate the fifteen most and fifteen least expressed samples corresponding to the edge. To select the “most” and “least” expressed samples, we assign each sample a summary “expression score.” The expression score is based on the expression values of the genes (limited to the top twenty genes with an odds ratio above 1) annotated to one or both of the pathways. Here, we show the heatmap of least expressed samples specific to the [Phosphate transport - Type II general secretion] relationship. **(c)** Clicking on a square in the heatmap directs a user to an experiment page (https://goo.gl/KYNhwB) based on the sample corresponding to that square. A user can use the experiment page to identify whether the expression values of genes specific to an edge and a selected sample differ from those recorded in other samples of the experiment. In this experiment page, the first three samples (labeled in black) are *P. aeruginosa* “baseline” replicates grown for 72 h in drop-flow biofilm reactors. The following three samples (labeled in blue) are *P. aeruginosa* grown for an additional 12 h (84 h total). Labels in blue indicate that the three 84 h replicates are in the heatmap of least expressed samples displayed on the [Phosphate transport – Type II general secretion] edge page.

We created a web interface for deeper examination of relationships present in the pathway co-occurrence network. The details we included in an edge-specific page (1) highlight up to twenty genes—annotated to either of the two pathways in the edge—contained in features that also contain this edge, after controlling for the total number of features that contain each gene, and (2) display the expression levels of these genes in each of the fifteen samples where they were most and least expressed. The quantity of information (twenty genes, thirty samples total) we choose to include in an edge page is intentionally limited so that users can review it in a reasonable amount of time.

To implement the functionality in (1), we computed an odds ratio for every gene annotated to one or both pathways in the edge. The odds ratio measures how often we observe a feature enriched for both the given gene and the edge of interest relative to how often we would expect to see this occurrence. We calculate the proportion of observed cases and divide by the expected proportion--equivalent to the frequency of the edge appearing in the model’s features.

Let *k* be the number of features from which the PathCORE-T network was built. *k_G_* is the number of features that contain gene *G* (i.e. *G* is in *k_G_* features’ gene signatures), *k_E_* the number of features that contain edge *E* (i.e. the two pathways connected by *E* are overrepresented in *k_E_* features), and *k_G & E_* the number of features that contain both gene *G* and edge *E*. The odds ratio is computed as follows:

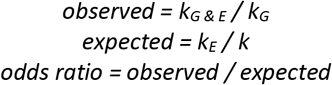

An odds ratio above 1 suggests that the gene is more likely to appear in features enriched for this pair of pathways. In the web interface, we sort the genes by their odds ratio to highlight genes most observed with the co-occurrence relationship.

The information specified in (2) requires an “expression score” for every sample. A sample expression score is calculated using the genes we selected in goal (1): it is the average of the normalized gene expression values weighted by the normalized gene odds ratio. Selection of the most and least expressed samples is based on these scores. We use two heatmaps to show the (maximum of twenty) genes’ expression values in each of the fifteen most and least expressed samples (Fig. 4b).

For each sample in an edge page, a user can examine how the expression values of the edge’s twenty genes in that sample compare to those recorded for all other samples in the dataset that are from the same experiment (Fig. 4c). Genes that display distinct expression patterns under a specific setting may be good candidates for follow-up studies.

## Results

### PathCORE-T software

Unsupervised methods can identify previously undiscovered patterns in large collections of data. PathCORE-T overlays curated knowledge after feature construction to help researchers interpret constructed features in the context of existing knowledgebases. Specifically, PathCORE-T aims to clarify how expert-annotated gene sets work together from a gene expression perspective. PathCORE-T starts from an unsupervised feature construction model. Before applying the software, users should evaluate models to make sure that they capture biological features in their dataset. Model evaluation can be performed in numerous ways depending on the setting and potential assessments include consistency across biological replicates, reconstruction error given a fixed dimensionality, and independent validation experiments. Tan et al. described several ways that models could be evaluated [7]. Datasets will vary in terms of their amenability to analysis by different model-building strategies, and researchers may wish to consult a recent review for more discussion of feature construction methods [33].

We implemented the methods contained in the PathCORE-T software in Python. The implementations of the primary steps are pip-installable (PyPI package name: PathCORE-T), and examples of how to use the methods in PathCORE-T are provided in the PathCORE-T-analysis repository (https://github.com/greenelab/PathCORE-T-analysis). We also implemented an optional step, which corrects for overlapping genes between pathway definitions, described by Donato et al. [10]. Though the algorithm had been described, no publicly available implementation existed. We provide this overlap correction algorithm as a Python package (PyPI package name: crosstalk-correction) available under the BSD 3-clause license. Each component of PathCORE-T can be used independently of each other (Fig. 1c).

Here, we present analyses that can be produced by applying the full PathCORE-T pipeline to models created from a transcriptomic compendium by an unsupervised feature construction algorithm. Input pathway definitions are “overlap-corrected” for each feature before enrichment analysis. An overlap-corrected, weighted pathway co-occurrence network is built by connecting the pairs of pathways that are overrepresented in features of the model. Finally, we remove edges that cannot be distinguished from a null model of random associations based on the results of a permutation test.

### Case study: *P. aeruginosa* eADAGE models annotated with KEGG pathways

We used PathCORE-T to create a network of co-occurring pathways out of the expression signatures extracted by eADAGE from a *P. aeruginosa* compendium [7]. For every feature, overlap correction was applied to the *P. aeruginosa* KEGG pathway annotations and overlap-corrected annotations were used in the overrepresentation analysis. PathCORE-T aggregates multiple networks by taking the union of the edges across all networks and summing the weights of common pathway-pathway connections. We do this to emphasize the co-occurrence relationships that are more stable [34]—that is, the relationships that appear across multiple models. Finally, we removed edges in the aggregate network that were not significant after FDR correction when compared to the background distributions generated from 10,000 permutations of the network. Used in this way, PathCORE-T software allowed for exploratory analysis of an existing well-validated model.

The eADAGE co-occurrence network that resulted from our exploratory analysis contained a number of pathway-pathway relationships that have been previously characterized by other means (Fig. 5). Three glucose catabolism processes co-occur in the network: glycolysis, pentose phosphate, and the Entner-Doudoroff pathway (Fig. 5a). We also found a cluster relating organophosphate and inorganic phosphate transport- and metabolism-related processes (Fig. 5b). Notably, phosphate uptake and acquisition genes were directly connected to the *hxc* genes that encode a type II secretion system. Studies in *P. aeruginosa* clinical isolates demonstrated that the Hxc secretion system was responsible for the secretion of alkaline phosphatases, which are phosphate scavenging enzymes [35,36] and the phosphate binding DING protein [37]. Furthermore, alkaline phosphatases, DING and the *hxc* genes are regulated by the transcription factor PhoB which is most active in response to phosphate limitation. The identification of this relationship by PathCORE-T as a global pattern suggested the role of type II secretion and phosphate limitation seen in a limited number of isolates may be generalizable to broader *P. aeruginosa* biology. As shown in Fig. 5c, we also identified linkages between two pathways involved in the catabolism of sulfur-containing molecules, taurine and methionine, and the general sulfur metabolism process. Other connections between pathways involved in the transport of iron (ferrienterobactin binding) [38] and zinc (the *znu* uptake system [39]) were identified (Fig. 5d). Interestingly, genes identified in the edge between the zinc transport and MacAB-TolC pathways included the *pvd* genes involved in pyoverdine biosynthesis and regulation, a putative periplasmic metal binding protein, as well as other components of an ABC transporter (genes PA2407, PA2408, and PA2409 at https://goo.gl/bfqOk8) [40]. PathCORE-T suggested a relationship between zinc and iron pathways in *P. aeruginosa* transcriptional data though such a relationship has not yet been described. Structural analysis of the iron-responsive regulator Fur found that it also productively binds zinc in *E. coli* and *Bacillus subtilis* providing a mechanism by which these pathways may be linked [41, 42].

**Figure 5.**
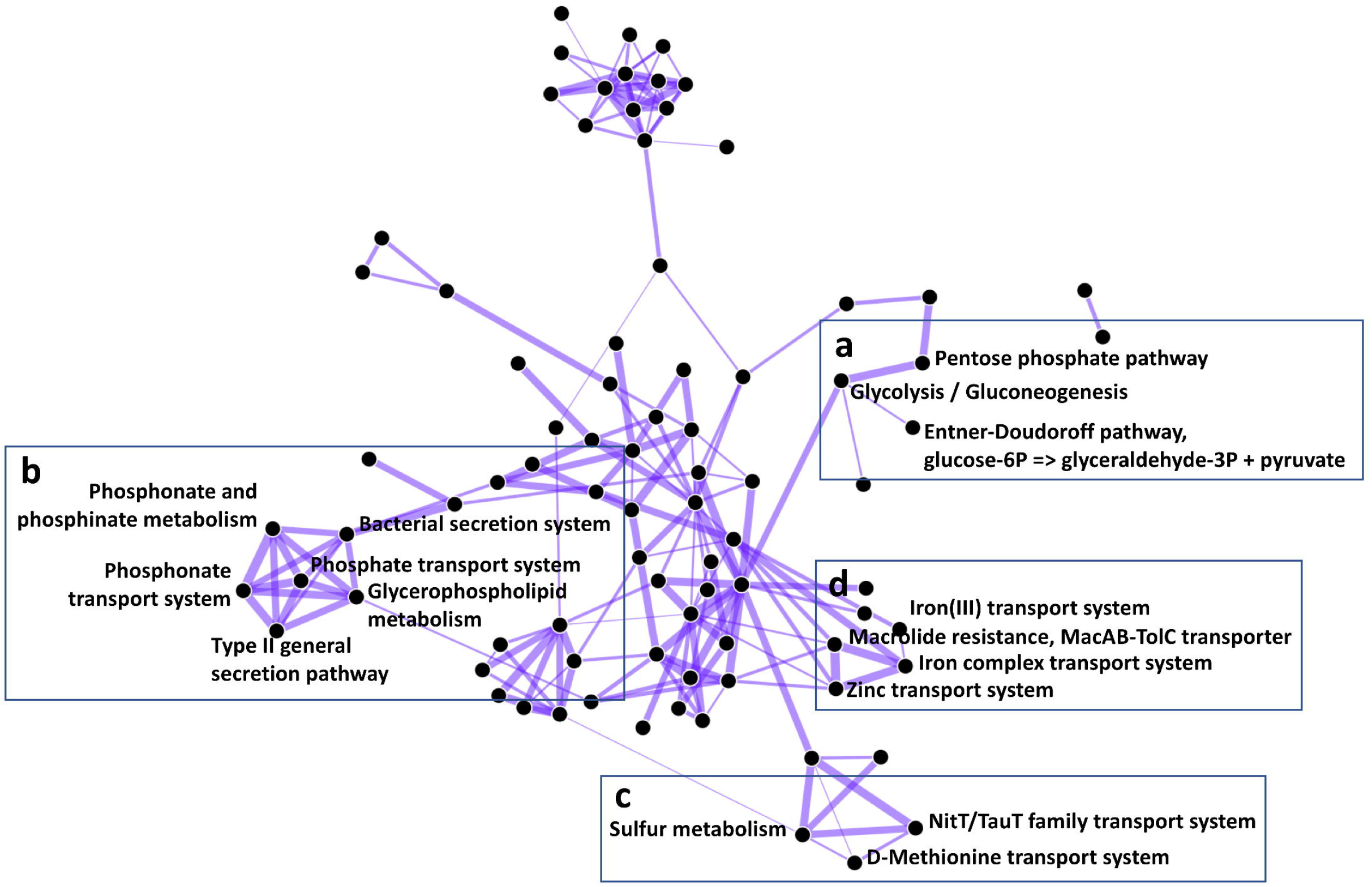
eADAGE features constructed from publicly available *P. aeruginosa* expression data describe known KEGG pathway-pathway relationships. (a) The glycerolipid metabolism, Entner-Doudoroff, glycolysis/gluconeogenesis, and pentose phosphate, pathways share common functions related to glucose catabolism. (b) Organophosphate and inorganic phosphate transport- and metabolism-related processes frequently co-occur with bacterial secretion systems. Here, we observe pairwise relationships between type II general secretion and phosphate-related processes. (c) Pathways involved in the catabolism of sulfur-containing molecules--taurine (NitT/TauT family transport) and methionine (D-Methionine transport), and the general sulfur metabolism process--are functionally linked. (d) The zinc transport, iron complex transport, and MacAB-TolC transporter systems are pairwise connected. The fully labeled network can be viewed at https://pathcore-demo.herokuapp.com/PAO1. The list of KEGG pathway-pathway relationships visualized in the network is available at the specified link (Ctrl + L for list view) and as a supplemental file for this paper.

The network constructed using the PathCORE-T framework had 203 edges between 89 pathways. For comparison, we constructed a KEGG pathway-pathway network where edges were drawn between pathways with significant gene overlap (FDR-corrected hypergeometric test < 0.05). The overlap-based network had 406 edges between 158 pathways. Only 35 of the edges in the PathCORE-T network were between pathways that shared genes, with an average Jaccard Index of only 0.035. The network constructed using PathCORE-T (with overlap-correction applied by default) captured pathway co-occurrences not driven by shared genes between pathways.

### Case study: TCGA’s pan-cancer compendium analyzed by NMF with PID pathways

PathCORE-T is not specific to a certain dataset, organism, or feature construction method. We constructed a 300-feature NMF model of TCGA pan-cancer gene expression data, which is comprised of 33 different cancer-types from various organ sites and applied the PathCORE-T software to those features. We chose NMF because it has been used in previous studies to identify biologically relevant patterns in transcriptomic data [12] and by many studies to derive molecular subtypes [43–45]. The 300 NMF features were analyzed using overlap-corrected PID pathways, a collection of 196 human cell signaling pathways with a particular focus on processes relevant to cancer [11].

PathCORE-T detected modules of co-occurring pathways that were consistent with our current understanding of cancer-related interactions (Fig. 6). Because cancer-relevant pathways were used, it was not surprising that cancer-relevant pathways appeared. However, the edges between those pathways were also encouraging. For example, a module composed of a FoxM1 transcription factor network, an E2F transcription factor network, Aurora B kinase signaling, ATR signaling, PLK1 signaling, and members of the Fanconi anemia DNA damage response pathway was densely connected (Fig 6a). The connections in this module recapitulated well known cancer hallmarks including cellular proliferation pathways and markers of genome instability, such as the activation of DNA damage response pathways [46]. We found that several pairwise pathway co-occurrences corresponded with previously reported pathway-pathway interactions [47–49]. We also observed a hub of pathways interacting with Wnt signaling, among them the regulation of nuclear Beta-catenin signaling, FGF signaling, and BMP signaling (Fig. 6b). The Wnt and BMP pathways are functionally integrated in biological processes that contribute to cancer progression [50]. Additionally, Wnt/Beta-catenin signaling is a well-studied regulatory system, and the effects of mutations in Wnt pathway components on this system have been linked to tumorigenesis [51]. Wnt/Beta-catenin and FGF together influence the directional migration of cancer cell clusters [52].

**Figure 6.**
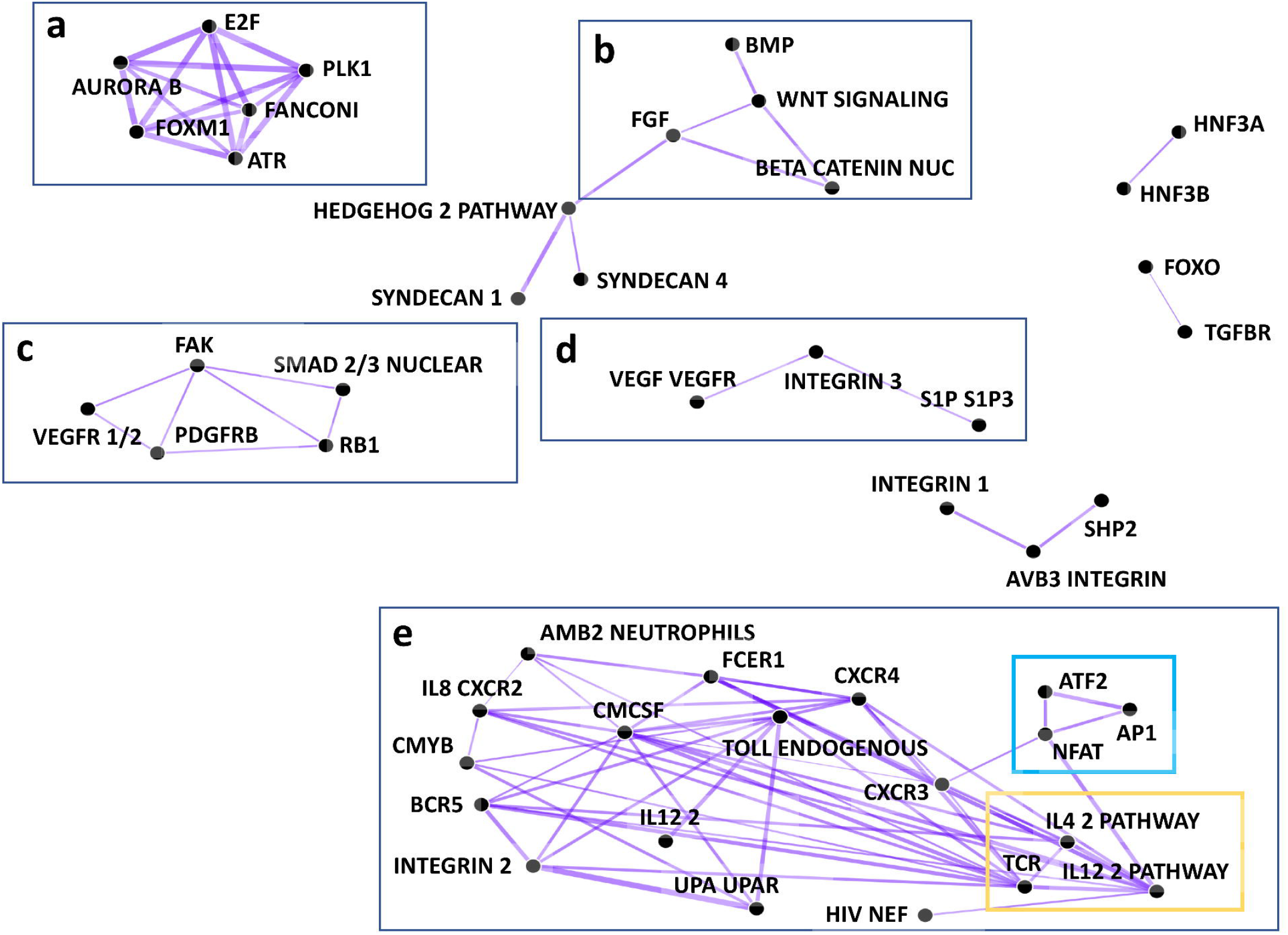
PID pathway-pathway relationships discovered in NMF features constructed from the TCGA pan-cancer gene expression dataset. (a) Pathways in this module are responsible for cell cycle progression. (b) Wnt signaling interactions with nuclear Beta-catenin signaling, FGF signaling, and BMP signaling have all been linked to cancer progression. (c) Here, we observe functional links between pathways responsible for angiogenesis and those responsible for cell proliferation. (d) The VEGF-VEGFR pathway interacts with the S1P3 pathway through Beta3 integrins. (e) This module contains many relationships related to immune system processes. The interaction cycle formed by T-Cell Receptor (TCR) signaling in naïve CD4+ T cells and IL-12/IL-4 mediated signaling events, outlined in yellow, is one well-known example. The cycle in blue is formed by the ATF2, NFAT, and AP1 pathways; pairwise co-occurrence of these three transcription factor networks may suggest that dysregulation of any one of these pathways can trigger variable oncogenic processes in the immune system. The list of PID pathway-pathway relationships visualized in the network is available as a supplemental file for this paper.

Two modules in the network related to angiogenesis, or the formation of new blood vessels (Fig. 6c, d). Tumors transmit signals that stimulate angiogenesis because a blood supply provides the necessary oxygen and nutrients for their growth and proliferation. One module relates angiogenesis factors to cell proliferation. This module connected the following pathways: PDGFR-beta signaling, FAK-mediated signaling events, VEGFR1 and VEGFR2-mediated signaling events, nuclear SMAD2/3 signaling regulation, and RB1 regulation (Fig. 6c). These functional connections are known to be involved in tumor proliferation [53–55]. The other module indicated a relationship by which the VEGF pathway interacts with the S1P3 pathway through Beta3 integrins (Fig. 6d). S1P3 is a known regulator of angiogenesis [56], and has been demonstrated to be associated with treatment-resistant breast cancer and poor survival [57]. Moreover, this interaction module has been observed to promote endothelial cell migration in human umbilical veins [58]. Taken together, this independent module may suggest a distinct angiogenesis process activated in more aggressive and metastatic tumors that is disrupted and regulated by alternative mechanisms [59].

Finally, PathCORE-T revealed a large, densely connected module of immune related pathways (Fig. 6e). We found that this module contains many co-occurrence relationships that align with immune system processes. One such example is the well characterized interaction cycle formed by T-Cell Receptor (TCR) signaling in naïve CD4+ T cells and IL-12/IL-4 mediated signaling events [60–62]. At the same time, PathCORE-T identifies additional immune-related relationships. We observed a cycle between the three transcription factor networks: ATF-2, AP-1, and CaN-regulated NFAT-dependent transcription. These pathways can take on different, often opposing, functions depending on the tissue and subcellular context. For example, ATF-2 can be an oncogene in one context (e.g. melanoma) and a tumor suppressor in another (e.g. breast cancer) [63]. AP-1, comprised of Jun/Fos proteins, is associated with both tumorigenesis and tumor suppression due to its roles in cell survival, proliferation, and cell death [64]. Moreover, NFAT in complex with AP-1 regulates immune cell differentiation, but dysregulation of NFAT signaling can lead to malignant growth and tumor metastasis [65]. The functional association observed between the ATF-2, AP-1, and NFAT cycle together within the immunity module might suggest that dysregulation within this cycle has profound consequences for immune cell processes and may trigger variable oncogenic processes.

Just as we did for the eADAGE-based *P. aeruginosa* KEGG pathways case study, we constructed a network only from PID pathways with significant gene overlap. The network constructed using PathCORE-T and NMF features had 119 edges between 57 pathways. The overlap-based network was much denser: it had 3826 edges between 196 pathways. This suggested a high degree of overlap between PID pathways. For the PathCORE-T NMF co-occurrence network, 96 of the edges were between pathways that shared genes. However, the average Jaccard Index for these pathway pairs remained low, at 0.058.

## Conclusions

Unsupervised analyses of genome-scale datasets that summarize key patterns in the data have the potential to improve our understanding of how a biological system operates via complex interactions between molecular processes. Feature construction algorithms capture coordinated changes in the expression of many genes as features. The genes that contribute most to each feature co-vary. However, interpreting the features generated by unsupervised approaches remains challenging. PathCORE-T creates a network of globally co-occurring pathways based on features created from the analysis of a compendium of gene expression data. Networks modeling the relationships between curated processes in a biological system offer a means for developing new hypotheses about which pathways influence each other and when. Our framework provides a data-driven characterization of the biological system at the pathway-level by identifying pairs of pathways that are overrepresented across many features.

PathCORE-T connects the features extracted from data to curated resources. It is important to note that PathCORE-T will only be able to identify pathways that occur in features of the underlying model, which means these pathways must be transcriptionally perturbed in at least some subset of the compendium. Models should be evaluated before analysis with PathCORE-T. The network resulting from PathCORE-T can help to identify global pathway-pathway relationships—a baseline network—that complements existing work to identify interactions between pathways in the context of a specific disease. The specific niche that PathCORE-T framework aims to fill is in revealing to researchers which gene sets are most closely related to each other in machine learning-based models of gene expression, which genes play a role in this co-occurrence, and which conditions drive this relationship.

**Project name**: PathCORE-T

**Project home page**: https://pathcore-demo.herokuapp.com

**Archived version**: https://github.com/greenelab/PathCORE-T-analysis/releases/tag/v1.2.0 (links to download .zip and .tar.gz files are provided here)

**Operating system**: Platform independent

**Programming language**: Python

**Other requirements**: Python 3 or higher

**License**: BSD 3-clause

## Declarations

### Ethics approval and consent to participate

Not applicable

### Consent for publication

Not applicable

### Availability of data and materials

Files for each of the PathCORE-T networks described in the results are provided in Supplementary material.

#### Data sets

*P. aeruginosa* eADAGE models: doi:10.5281/zenodo.583172

*P. aeruginosa* gene expression compendium: doi:10.5281/zenodo.583694

The normalized gene expression compendium provided in this Zenodo record contains datasets on the GPL84 from ArrayExpress as of 31 July 2015. It includes 1,051 samples grown in 78 distinct medium conditions, 128 distinct strains and isolates, and dozens of different environmental parameters [7]. Tan et al. compiled this dataset and used it to construct the 10 eADAGE models from which we generate the eADAGE-based *P. aeruginosa* KEGG network [7]. We use this same data compendium to generate the NMF *P. aeruginosa* model. The script used to download and process those datasets into the compendium is available at https://goo.gl/YjOEQl.

TCGA pan-cancer dataset: doi:10.5281/zenodo.56735

The pan-cancer dataset was compiled using data from all 33 TCGA cohorts. It was generated by the TCGA Research Network: http://cancergenome.nih.gov/ [14]. RNA-seq data was downloaded on 8 March 2016 from UCSC Xena (https://xenabrowser.net/datapages/). Gene expression was measured using the Illumina HiSeq technology. More information and the latest version of the dataset can be found at https://xenabrowser.net/datapages/?dataset=TCGA.PANCAN.sampleMap/HiSeqV2&host=https://tcga.xenahubs.net.

#### Source code

<u>PathCORE-T analysis</u>: (https://github.com/greenelab/PathCORE-T-analysis/tree/v1.2.0) This repository contains all the scripts to reproduce the analyses described in this paper. The Python scripts here should be used as a starting point for new PathCORE-T analyses. Instructions for setting up a web application for a user’s specific PathCORE-T analysis are provided in this repository’s README.

<u>Overlap correction</u>: (https://github.com/kathyxchen/crosstalk-correction/tree/v1.0.4) Donato et. al’s procedure for overlap correction [10] is a pip-installable Python package ‘crosstalk-correction’ that is separate from, but listed as a dependency in, PathCORE-T. It is implemented using NumPy [31].

<u>PathCORE-T methods</u>: (https://github.com/greenelab/PathCORE-T/tree/v1.0.2) The methods included in the PathCORE-T analysis workflow (Fig. 4c) are provided as a pip-installable Python package ‘pathcore’. It is implemented using Pandas [66], SciPy (specifically, scipy.stats) [67], StatsModels [68], and the crosstalk-correction package.

<u>PathCORE-T demo application</u>: (https://github.com/kathyxchen/PathCORE-T-demo/tree/v2.1) The project home page, https://pathcore-demo.herokuapp.com provides links to

1. The web application for the eADAGE-based KEGG *P. aeruginosa* described in the first case study.
2. A view of the NMF-based PID pathway co-occurrence network described in the second case study.
3. A quick view page where users can temporarily load and visualize their own network file (generated from the PathCORE-T analysis).

## Authors’ contributions

KMC implemented the software, performed the analyses, and drafted the manuscript. JT and GPW contributed computational reagents. KMC, DAH, and CSG designed the project. KMC, JT, GPW, GD, DAH, and CSG interpreted the results. JT, GPW, GD, DAH, and CSG provided critical feedback and revisions on the manuscript. JT and GPW reviewed source code.

## Acknowledgements

The authors would also like to thank Daniel Himmelstein, Dongbo Hu, Kurt Wheeler, and René Zelaya for helping to review the source code.

